## Supplementary Information for "A doubly stochastic renewal framework for partitioning spiking variability"

February 21, 2024

### 1 Supplementary Notes

#### 1.1 Derivations of mathematical results for renewal point processes

**Definition 1** (Ordinary and equilibrium point processes)**.** A point process can be called ordinary or equilibrium depending on the time at which we set the origin $t = 0$. The point process is called ordinary if $t = 0$ is set at the time of the first spike. For an ordinary renewal point process, all interspike intervals (ISIs) are i.i.d. with the probability density $f(x)$. If an ordinary point process runs for a long time and then the time origin is picked randomly, then the resulting point process is called equilibrium [1]. This process is in a steady state because the time at which the process started is far from the time $t = 0$ at which we start sampling the process. For an equilibrium renewal point process, the interval from $t = 0$ to the first spike time $t_1$ has a different probability density $f_1(x)$ than all subsequent ISIs.

**Theorem 1.** Consider an equilibrium renewal point process defined by the ISI probability density $f(x)$, meaning that after generating a spike the probability that the next spike happens in the interval $[x, x + dx]$ is $f(x)dx$. Denoting the first three central moments of $f(x)$ by $\mu$, $\sigma^2$, and $\mu_3$, the mean $\mathrm{E}(\boldsymbol{N}_T)$ and the variance $\mathrm{Var}(\boldsymbol{N}_T)$ of the spike count $\boldsymbol{N}_T$ in a bin with the size $T$ are

$$\begin{aligned}
\mathrm{E}(\boldsymbol{N}_T) &= \frac{T}{\mu}, \\
\mathrm{Var}(\boldsymbol{N}_T) &= \frac{\sigma^2}{\mu^3}T + \frac{\sigma^4}{2\mu^4} + \frac{1}{6} - \frac{\mu_3}{3\mu^3} + \mathcal{O}(T^{-1}).
\end{aligned}$$

*Proof.* Let us denote the time of the $r^{\text{th}}$ spike by $x$, its probability density function by $k_r(x)$, and its cumulative distribution function by $K_r(x)$. Let us define the functions:

$$\begin{aligned}
H(T) &:= E(\boldsymbol{N}_T), \\
\psi(T) &:= E(\boldsymbol{N}_T(\boldsymbol{N}_T + 1)).
\end{aligned}$$

With these definition, the variance of the spike count can be written as

$$\mathrm{Var}(\boldsymbol{N}_T) = \psi(T) - H(T) - H^2(T). \tag{1}$$

We next express $H(T)$ and $\psi(T)$ in terms of the density function $f(x)$ and its moments.

$$\begin{aligned}
H(T) &= \sum_{r=0}^{\infty} r\mathbb{P}(\boldsymbol{N}_T = r) = \sum_{r=0}^{\infty} r(K_r(T) - K_{r+1}(T)) \\
&= \sum_{r=0}^{\infty} rK_r(T) - \sum_{r=0}^{\infty}(r+1)K_{r+1}(T) + \sum_{r=0}^{\infty} K_{r+1}(T) \\
&= \sum_{r=1}^{\infty} rK_r(T) - \sum_{r=1}^{\infty} rK_r(T) + \sum_{r=1}^{\infty} K_r(T) \\
&= \sum_{r=1}^{\infty} K_r(T).
\end{aligned}$$

Similarly, we have

$$\begin{aligned}
\psi(T) &= \sum_{r=0}^{\infty} r(r+1)\mathbb{P}(\boldsymbol{N}_T = r) = \sum_{r=1}^{\infty} r(r+1)(K_r(T) - K_{r+1}(T)) \\
&= \sum_{r=1}^{\infty} r(r+1)K_r(T) - \sum_{r=1}^{\infty}(r+1)(r+2)K_{r+1}(T) + \sum_{r=1}^{\infty} 2(r+1)K_{r+1}(T) \\
&= \sum_{r=1}^{\infty} r(r+1)K_r(T) - \sum_{r=2}^{\infty} r(r+1)K_r(T) + \sum_{r=2}^{\infty} 2rK_r(T) \\
&= 2K_1(T) + \sum_{r=2}^{\infty} 2rK_r(T) = \sum_{r=1}^{\infty} 2rK_r(T)\ .
\end{aligned}$$

Using these two equalities, we write the Laplace transform of $\psi$ and $H$, $\psi^*(s) = \mathcal{L}\{\psi\}(s)$ and $H^*(s) = \mathcal{L}\{H\}(s)$, in terms of the Laplace transform of $k_r$:

$$H^*(s) = \sum_{r=1}^{\infty} K_r^*(s) = \sum_{r=1}^{\infty} \frac{1}{s} k_r^*(s), \tag{2}$$

$$\psi^*(s) = \sum_{r=1}^{\infty} 2r K_r^*(s) = \sum_{r=1}^{\infty} \frac{2}{s} r k_r^*(s). \tag{3}$$

Let us denote the time of the first spike $t_1$, the interspike interval between the 1st and 2nd spikes by $t_2$, and similarly the time between $(r-1)^{\text{th}}$ and $r^{\text{th}}$ spikes by $t_r$. Recalling that $x$ is the time of the $r^{\text{th}}$ spike, we have $x = \sum_{i=1}^{r} t_i$. Thus,

$$k_r(x) = (f_1 * \overbrace{f * \cdots * f}^{r-1})(x), \tag{4}$$

where $f_1(.)$ is the probability density of the first interval $t_1$, and $f(.)$ is the probability density for the subsequent $r-1$ ISIs $t_i$. Since we have an equilibrium point process, $f_1$ and $f$ are different functions. The operator $*$ denotes the time-domain convolution. Applying the Laplace transform to Eq. 4, we obtain

$$k_r^*(s) = f_1^*(s)(f^*(s))^{r-1}. \tag{5}$$

One can show that we have [1, 2]

$$f_1^*(s) = \frac{1 - f^*(s)}{\mu s}. \tag{6}$$

Substituting Eq. 5 into Eqs. 2,3 and using Eq. 6, we obtain

$$H^*(s) = \frac{1}{\mu s^2}, \tag{7}$$

$$\psi^*(s) = \frac{2}{\mu s^2 (1 - f^*(s))}. \tag{8}$$

Taking the inverse Laplace transform in Eq. 7, we find

$$H(T) = \frac{T}{\mu}. \tag{9}$$

For a general case, we can not obtain a closed form for the inverse Laplace transform in Eq. 8 to calculate $\psi(T)$ exactly, but we can approximate it. If there exist $s_0 > 0$ such that for $Re(s) > -s_0$, $f^*(s)$ is analytical, then we can expand the Laplace transform $f^*(s)$ in terms of moments $m_n$ of the probability density $f(x)$ [3]:

$$f^*(s) = \sum_{n=0}^{\infty} \frac{m_n}{n!}(-s)^n \approx 1 - \mu s + \frac{m_2}{2} s^2 - \frac{m_3}{3!} s^3.$$

Here $m_2 = \sigma^2 + \mu^2$ and $m_3 = \mu_3 + 3\mu\sigma^2 + \mu^3$ where $\mu_3 = E((x-\mu)^3)$. Now using Eq. 8, we can estimate $\psi^*(s)$:

$$\psi^*(s) \approx \frac{2}{\mu s^2 (\mu s - \frac{m_2}{2} s^2 + \frac{m_3}{3!} s^3)} .$$

We expand this function around $s = 0$ and use $(1-x)^{-1} = 1 + x + x^2 + x^3 + \cdots$ to obtain

$$\begin{aligned}\psi^*(s) &\approx \frac{2}{\mu^2 s^3} \frac{1}{(1 - \frac{m_2}{2\mu}s + \frac{m_3}{3!\mu}s^2)} \\ &\approx \frac{2}{\mu^2 s^3}\left(1 + \frac{m_2}{2\mu}s - \frac{m_3}{3!\mu}s^2 + \left(\frac{m_2}{2\mu}\right)^2 s^2 + \mathcal{O}(s^3)\right).\end{aligned}$$

Now we express $m_2$ and $m_3$ in terms of $\mu$, $\sigma$ and $\mu_3$ and get

$$\begin{aligned}\psi^*(s) &\approx \frac{2}{\mu^2 s^3}\left(1 + \frac{\sigma^2+\mu^2}{2\mu}s + \left(\left(\frac{\sigma^2+\mu^2}{2\mu}\right)^2 - \frac{\mu^3 + 3\mu\sigma^2 + \mu_3}{6\mu}\right)s^2 + \mathcal{O}(s^3)\right) \\ &= \frac{2}{\mu^2 s^3}\left(1 + \frac{\sigma^2+\mu^2}{2\mu}s + \left(\frac{\sigma^4}{4\mu^2} + \frac{\mu^2}{12} - \frac{\mu_3}{6\mu}\right)s^2 + \mathcal{O}(s^3)\right) \\ &= \frac{2}{\mu^2}\frac{1}{s^3} + \frac{\sigma^2+\mu^2}{\mu^3}\frac{1}{s^2} + \left(\frac{\sigma^4}{2\mu^4} + \frac{1}{6} - \frac{\mu_3}{3\mu^3}\right)\frac{1}{s} + \mathcal{O}(1)\ .\end{aligned}$$

Taking the inverse Laplace transform, we have

$$\psi(T) = \frac{1}{\mu^2}T^2 + \frac{\sigma^2+\mu^2}{\mu^3}T + \frac{\sigma^4}{2\mu^4} + \frac{1}{6} - \frac{\mu_3}{3\mu^3} + \mathcal{O}(T^{-1}). \tag{10}$$

Using Eqs. 1,9,10, we obtain the final result

$$\begin{aligned}\mathrm{Var}(\boldsymbol{N}_T) &= \frac{1}{\mu^2}T^2 + \frac{\sigma^2+\mu^2}{\mu^3}T + \frac{\sigma^4}{2\mu^4} + \frac{1}{6} - \frac{\mu_3}{3\mu^3} - \frac{1}{\mu}T - \frac{1}{\mu^2}T^2 + \mathcal{O}(T^{-1}) \\ &= \frac{\sigma^2}{\mu^3}T + \frac{\sigma^4}{2\mu^4} + \frac{1}{6} - \frac{\mu_3}{3\mu^3} + \mathcal{O}(T^{-1}).\end{aligned}$$

□

**Theorem 2.** For an equilibrium doubly stochastic renewal point process $\{g(\cdot), \boldsymbol{\lambda}(t)\}$ with a stationary firing rate $\boldsymbol{\lambda}(t)$ changing on a timescale $\tau$ longer than the bin size $\tau > T$, we have $\mathrm{E}[\boldsymbol{N}_T|\boldsymbol{\lambda}] = \boldsymbol{\lambda}T$.

*Proof.* Since $\boldsymbol{\lambda}(t)$ changes on a timescale longer than $T$, we can consider $\boldsymbol{\lambda}$ to be approximately constant within a bin. Then, within a single bin, the spike generating process is a renewal point process fully defined by its ISI probability density, which we denote by $\boldsymbol{f_\lambda}$ since it depends on the value of $\boldsymbol{\lambda}$ in the bin. Theorem 1 asserts that for such a renewal process, $E(\boldsymbol{N}_T) = T/\mu$, where $\mu$ is first moment of the ISI probability density. Hence, $\mathrm{E}[\boldsymbol{N}_T|\boldsymbol{\lambda}] = T/\boldsymbol{\mu_{f_\lambda}}$, where $\boldsymbol{\mu_{f_\lambda}}$ is the mean of the probability density $\boldsymbol{f_\lambda}$. Since $\boldsymbol{\lambda}$ is constant, by the definition of the renewal process, the density $\boldsymbol{f_\lambda}$ is a linear transformation of $g(\cdot)$, hence the mean of $\boldsymbol{f_\lambda}$ is

$$\boldsymbol{\mu_{f_\lambda}} = \frac{\mu_g}{\boldsymbol{\lambda}} = \frac{1}{\boldsymbol{\lambda}}.$$

As a result, $\mathrm{E}[\boldsymbol{N}_T|\boldsymbol{\lambda}] = T/(1/\boldsymbol{\lambda}) = \boldsymbol{\lambda}T$.

□

**Theorem 3.** For an equilibrium doubly stochastic renewal point process $\{g(\cdot), \boldsymbol{\lambda}(t)\}$ with a stationary firing rate $\boldsymbol{\lambda}(t)$ changing on a timescale $\tau$ longer than the bin size $\tau > T$, we have

$$\mathrm{E}[\mathrm{Var}(\boldsymbol{N}_T|\boldsymbol{\lambda})] = \left(\frac{\sigma_g}{\mu_g}\right)^2 \mathrm{E}[\boldsymbol{N}_T] + \frac{1}{6} + \frac{1}{2}\left(\frac{\sigma_g}{\mu_g}\right)^4 - \frac{1}{3}\frac{\mu_{3_g}}{\mu_g^3} + \mathcal{O}(T^{-1}).$$

*Proof.* Since $\boldsymbol{\lambda}(t)$ changes on a timescale longer than $T$, we can consider $\boldsymbol{\lambda}$ to be approximately constant within a bin. The proof of Theorem 1 shows that we can expand $\mathrm{Var}(\boldsymbol{N_T})$ in terms of powers of the bin size $T$ where the powers go from $-\infty$ to 1. Since within a single bin, the spike generating process is a renewal point process defined by its ISI density, we can use the result of Theorem 1 to write

$$\mathrm{Var}(\boldsymbol{N}_T|\boldsymbol{\lambda}) = \sum_{i=-\infty}^{1} \boldsymbol{c_i} T^i.$$

Here $\boldsymbol{c_i}$ are random variables that depend on $\boldsymbol{f_\lambda}$.

For moderately large bin size $T \gg 1/\mathrm{E}[\boldsymbol{\lambda}]$, we can approximate $\mathrm{Var}(\boldsymbol{N}_T|\boldsymbol{\lambda})$ by the two largest powers of $T$. The proof of Theorem 1 shows that the coefficients $\boldsymbol{c_0}$ and $\boldsymbol{c_1}$ are functions of the mean, standard deviation and third central moment of the probability density $\boldsymbol{f_\lambda}$, which we denote by $\boldsymbol{\mu_{f_\lambda}}$, $\boldsymbol{\sigma_{f_\lambda}}$ and $\boldsymbol{\mu_{3_{f_\lambda}}}$, respectively. Thus, using the result of Theorem 1, we can write

$$\mathrm{Var}(\boldsymbol{N}_T|\boldsymbol{\lambda}) = \frac{\boldsymbol{\sigma}^2_{\boldsymbol{f_\lambda}}}{\boldsymbol{\mu}^3_{\boldsymbol{f_\lambda}}} T + \frac{\boldsymbol{\sigma}^4_{\boldsymbol{f_\lambda}}}{2\boldsymbol{\mu}^4_{\boldsymbol{f_\lambda}}} + \frac{1}{6} - \frac{\boldsymbol{\mu}_{3\boldsymbol{f_\lambda}}}{3\boldsymbol{\mu}^3_{\boldsymbol{f_\lambda}}} + \mathcal{O}(T^{-1}).$$

By the statement of Theorem 2, we have $\mathrm{E}[\boldsymbol{N}_T|\boldsymbol{\lambda}] = \boldsymbol{\lambda}T = T/\boldsymbol{\mu_{f_\lambda}}$, from which we express $T$ as $T = \boldsymbol{\mu_{f_\lambda}}\mathrm{E}[\boldsymbol{N}_T|\boldsymbol{\lambda}]$ and thus we rewrite the equation for $\mathrm{Var}(\boldsymbol{N}_T|\boldsymbol{\lambda})$:

$$\mathrm{Var}(\boldsymbol{N}_T|\boldsymbol{\lambda}) = \left(\frac{\boldsymbol{\sigma_{f_\lambda}}}{\boldsymbol{\mu_{f_\lambda}}}\right)^2 \mathrm{E}[\boldsymbol{N}_T|\boldsymbol{\lambda}] + \frac{1}{6} + \frac{1}{2}\left(\frac{\boldsymbol{\sigma_{f_\lambda}}}{\boldsymbol{\mu_{f_\lambda}}}\right)^4 - \frac{1}{3}\frac{\boldsymbol{\mu}_{3\boldsymbol{f_\lambda}}}{\boldsymbol{\mu}^3_{\boldsymbol{f_\lambda}}} + \mathcal{O}(T^{-1}).$$

Since $\boldsymbol{\lambda}$ is constant within a bin, by the definition of a renewal process, the density $\boldsymbol{f_\lambda}$ is a linear transformation of $g(\cdot)$, hence we can express the moments of $\boldsymbol{f_\lambda}$ in terms of the moments of $g(\cdot)$:

$$\boldsymbol{\mu_{f_\lambda}} = \frac{\mu_g}{\boldsymbol{\lambda}} = \frac{1}{\boldsymbol{\lambda}}, \quad \boldsymbol{\sigma}^2_{\boldsymbol{f_\lambda}} = \frac{\sigma_g^2}{\boldsymbol{\lambda}^2}, \quad \boldsymbol{\mu}_{3\boldsymbol{f_\lambda}} = \frac{\mu_{3_g}}{\boldsymbol{\lambda}^3}.$$

In these relations, $\boldsymbol{\mu_{f_\lambda}}$, $\boldsymbol{\sigma_{f_\lambda}}$ and $\boldsymbol{\mu_{3_{f_\lambda}}}$ are random variables that fluctuate with $\boldsymbol{\lambda}$, which changes stochastically across time bins, whereas $\mu_g$, $\sigma_g$ and $\mu_{3_g}$ are fixed because the function $g(\cdot)$ is fixed. Using these relations, we arrive at

$$\mathrm{Var}(\boldsymbol{N}_T|\boldsymbol{\lambda}) = \left(\frac{\sigma_g}{\mu_g}\right)^2 \mathrm{E}[\boldsymbol{N}_T|\boldsymbol{\lambda}] + \frac{1}{6} + \frac{1}{2}\left(\frac{\sigma_g}{\mu_g}\right)^4 - \frac{1}{3}\frac{\mu_{3_g}}{\mu_g^3} + \mathcal{O}(T^{-1}).$$

For pair of random variables $\boldsymbol{X}$ and $\boldsymbol{Y}$, the Law of Total Expectation (LOTE) [4] states that $E(\boldsymbol{Y}) = \mathrm{E}[\mathrm{E}(\boldsymbol{Y}|\boldsymbol{X})]$. Taking the expectation over $\boldsymbol{\lambda}$ and applying LOTE in the previous equation, we obtain the final result

$$\mathrm{E}[\mathrm{Var}(\boldsymbol{N}_T|\boldsymbol{\lambda})] = \left(\frac{\sigma_g}{\mu_g}\right)^2 \mathrm{E}[\boldsymbol{N}_T] + \frac{1}{6} + \frac{1}{2}\left(\frac{\sigma_g}{\mu_g}\right)^4 - \frac{1}{3}\frac{\mu_{3_g}}{\mu_g^3} + \mathcal{O}(T^{-1}).$$

□

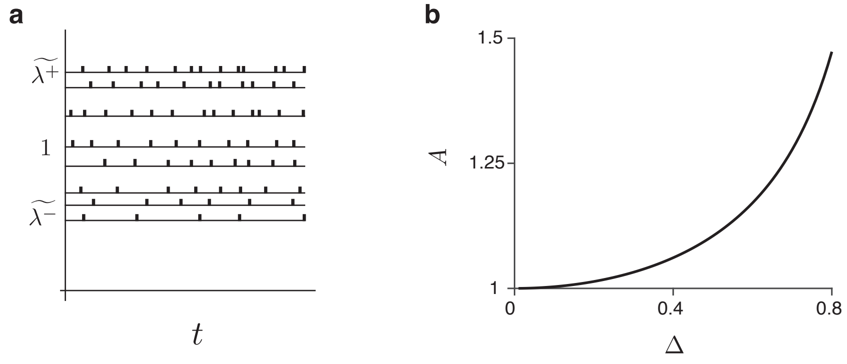

**Supplementary Figure 1.** Estimation error of the deterministic time rescaling method. **a**, We consider the case in which the firing rate is constant over time within a trial and fluctuates from trial to trial sampled from a uniform distribution over the interval $[\lambda^-, \lambda^+]$ Hz. Since the DTR method makes the assumption that the variance of the firing rate is zero, it mixes the ISIs from different trials with different firing rates. Intuitively, this mixing increases the estimated variance of ISIs. **b**, The estimated $\phi_{\mathrm{DTR}}$ is linearly related to the ground-truth $\phi$ via $\phi_{\mathrm{DTR}} = A\phi + (A-1)$. The parameter $A$ is a function of $\Delta = (\lambda^+ - \lambda^-)/(2\lambda_0)$ which controls the trial-to-trial variance of the firing rate.

#### 1.2 Bias of the Deterministic Time Rescaling method

To estimate the bias of the Deterministic Time Rescaling (DTR) method, we consider a doubly stochastic renewal point process $\{g(\cdot), \boldsymbol{\lambda}(t)\}$ in a particular time bin $[t, t+T]$. We assume that the firing rate is constant within a single bin but fluctuates from trial to trial with a uniform distribution over the interval $[\lambda^-, \lambda^+]$.

The DTR method assumes that the firing rate is deterministic and estimates the firing rate simply as the average spike count in a bin across trials. In our setting, this estimate produces

$$\lambda_0 := \frac{1}{T}\mathrm{E}(N) = \frac{1}{T}\mathrm{E}(\mathrm{E}(N|\lambda)) = \frac{1}{T}\mathrm{E}(\lambda T) = \mathrm{E}(\lambda) = \frac{\lambda^- + \lambda^+}{2}.$$

Using this estimated firing rate $\lambda_0$, the DTR method then rescales ISIs to map the spikes from real to operational time. This rescaling maps the highest firing rate $\lambda^+$ to $\widetilde{\lambda^+} = \lambda^+/\lambda_0 > 1$, and the lowest firing rate $\lambda^-$ to $\widetilde{\lambda^-} = \lambda^-/\lambda_0 < 1$, and $\lambda_0$ maps to $\widetilde{\lambda_0} = 1$ Hz. Finally, the DTR method estimates $\phi$ as $\mathrm{CV}^2$ of the rescaled ISIs by estimating the mean $\mu_n$ and standard deviation $\sigma_n$ of the rescaled ISIs. To calculate what this estimate will produce, we need to calculate the probability density $f(\tau)$ of rescaled ISIs. Rescaling time with the DTR method mixes ISIs from different trials, assuming they all came from the same firing rate, into a single $f(\tau)$ (Supplementary Fig. 1a). Thus, we have

$$f(\tau) = \int_{\widetilde{\lambda^-}}^{\widetilde{\lambda^+}} f(\tau|\lambda)\mathbb{P}(\lambda)d\lambda.$$

Since within a bin, the only contribution of $\lambda$ is to scale the ISIs, the conditional distribution is $f(\tau|\lambda) = \lambda g(\lambda\tau)$. Since we assumed a uniform distribution for the firing rate across trials,

$\mathbb{P}(\lambda) = \lambda/(\widetilde{\lambda^+} - \widetilde{\lambda^-})$ which leads to

$$f(\tau) = \frac{1}{\widetilde{\lambda^+} - \widetilde{\lambda^-}} \cdot \int_{\widetilde{\lambda^-}}^{\widetilde{\lambda^+}} \lambda g(\lambda\tau) d\lambda.$$

In theory, the first and second moments of the ISI probability density $g(\cdot)$ are

$$\begin{aligned} \mu_0 &= \int_0^\infty \tau g(\tau) d\tau = 1, \\ \sigma_0^2 &= \int_0^\infty (\tau - \mu_0)^2 g(\tau) d\tau = \phi. \end{aligned}$$

Using the DTR method, then mean $\mu_n$ of the rescaled ISIs is

$$\begin{aligned} \mu_n &= \int_0^\infty \tau f(\tau) d\tau = \int_0^\infty d\tau\tau \int_{\widetilde{\lambda^-}}^{\widetilde{\lambda^+}} f(\tau|\lambda)\mathbb{P}(\lambda) d\lambda \\ &= \frac{1}{\widetilde{\lambda^+} - \widetilde{\lambda^-}} \int_0^\infty d\tau\tau \int_{\widetilde{\lambda^-}}^{\widetilde{\lambda^+}} \lambda g(\lambda\tau) d\lambda = \frac{1}{\widetilde{\lambda^+} - \widetilde{\lambda^-}} \int_{\widetilde{\lambda^-}}^{\widetilde{\lambda^+}} d\lambda\lambda \int_0^\infty \tau g(\lambda\tau) d\tau \\ &= \frac{1}{\widetilde{\lambda^+} - \widetilde{\lambda^-}} \int_{\widetilde{\lambda^-}}^{\widetilde{\lambda^+}} d\lambda\lambda \int_0^\infty \frac{\tau'}{\lambda} g(\tau') \frac{d\tau'}{\lambda} = \frac{1}{\widetilde{\lambda^+} - \widetilde{\lambda^-}} \int_{\widetilde{\lambda^-}}^{\widetilde{\lambda^+}} \frac{1}{\lambda} d\lambda \int_0^\infty \tau' g(\tau') d\tau' = \frac{\ln(\widetilde{\lambda^+}/\widetilde{\lambda^-})}{\widetilde{\lambda^+} - \widetilde{\lambda^-}}. \end{aligned}$$

We see that the estimated mean ISI $\mu_n$ is not equal 1 s despite the fact that the linear average of the firing rate is $\widetilde{\lambda}_0 = 1$ Hz. This result shows that neglecting fluctuations in the firing rate and replacing it with a deterministic firing rate will produce the estimated mean ISI that is not necessarily consistent with the average firing rate. If trial-to-trial fluctuations of the firing rate are small compared to the average firing rate: $\widetilde{\lambda^+} - \widetilde{\lambda^-} \ll \widetilde{\lambda}_0 = 1$, then we recover $\mu_n \approx 1$ s. The equality only holds in the limit of zero variance, that is, for a deterministic firing rate.

For the standard deviation of the rescaled ISIs $\sigma_n$, we have

$$\begin{aligned} \widetilde{\sigma}_n^2 &= \int_0^\infty (\tau - \widetilde{\mu}_n)^2 f(\tau) d\tau = \int_0^\infty (\tau - \widetilde{\mu}_n)^2 \int_{\widetilde{\lambda^-}}^{\widetilde{\lambda^+}} f(\tau|\lambda)\mathbb{P}(\lambda) d\lambda d\tau \\ &= \frac{1}{\widetilde{\lambda^+} - \widetilde{\lambda^-}} \int_{\widetilde{\lambda^-}}^{\widetilde{\lambda^+}} d\lambda\lambda \int_0^\infty (\tau - \widetilde{\mu}_n)^2 g(\lambda\tau) d\tau \\ &= \frac{1}{\widetilde{\lambda^+} - \widetilde{\lambda^-}} \int_{\widetilde{\lambda^-}}^{\widetilde{\lambda^+}} d\lambda\lambda \int_0^\infty \left(\frac{\tau'}{\lambda} - \widetilde{\mu}_n\right)^2 g(\tau') \frac{d\tau'}{\lambda} \\ &= \frac{1}{\widetilde{\lambda^+} - \widetilde{\lambda^-}} \int_{\widetilde{\lambda^-}}^{\widetilde{\lambda^+}} d\lambda \int_0^\infty \left(\frac{\tau'^2}{\lambda^2} + \widetilde{\mu}_n^2 - 2\frac{\tau'}{\lambda}\widetilde{\mu}_n\right) g(\tau') d\tau' \\ &= \frac{1}{\widetilde{\lambda^+} - \widetilde{\lambda^-}} \int_{\widetilde{\lambda^-}}^{\widetilde{\lambda^+}} \left(\frac{1+\phi}{\lambda^2} + \widetilde{\mu}_n^2 - 2\frac{\widetilde{\mu}_n}{\lambda}\right) d\lambda \\ &= \frac{1}{\widetilde{\lambda^+} - \widetilde{\lambda^-}} \left((1+\phi)\left(\frac{1}{\widetilde{\lambda^-}} - \frac{1}{\widetilde{\lambda^+}}\right) + \widetilde{\mu}_n^2(\widetilde{\lambda^+} - \widetilde{\lambda^-}) - 2\widetilde{\mu}_n \ln\left(\frac{\widetilde{\lambda^+}}{\widetilde{\lambda^-}}\right)\right) \\ &= \frac{(1+\phi)}{\widetilde{\lambda^+}\widetilde{\lambda^-}} + \widetilde{\mu}_n^2 - 2\widetilde{\mu}_n^2 = \frac{(1+\phi)}{\widetilde{\lambda^+}\widetilde{\lambda^-}} - \widetilde{\mu}_n^2 \ . \end{aligned}$$

Then the estimated $\phi_{\mathrm{DTR}}$ is

$$\phi_{\mathrm{DTR}} = \frac{\widetilde{\sigma}_n^2}{\widetilde{\mu}_n^2} = A\phi + (A-1),$$

where

$$A = \frac{(\widetilde{\lambda^+} - \widetilde{\lambda^-})^2}{\widetilde{\lambda^+}\widetilde{\lambda^-}(\ln(\widetilde{\lambda^+}/\widetilde{\lambda^-}))^2} = \frac{4\Delta^2}{(1-\Delta^2)(\ln(\frac{1+\Delta}{1-\Delta}))^2}.$$

Here we defined $\Delta = \frac{\lambda^+ - \lambda^-}{2\lambda_0} = \frac{\lambda^+ - \lambda^-}{\lambda^+ + \lambda^-}$, which is a control parameter that indicates how much the firing rate process deviates from the deterministic assumption. $\Delta = 0$ corresponds to the deterministic firing rate which leads to $A = 1$ and we recover $\phi_{\mathrm{DTR}} = \phi$. The dependence of $A$ on $\Delta$ is a monotonically increasing function (Supplementary Fig. 1b), hence increasing $\Delta$ leads to a larger error in $\phi_{\mathrm{DTR}}$.

#### 1.3 Biases of the Minimum Ratio method

The Minimum Ratio (MR) method for partitioning spiking variability [5] starts with an assumption

$$\mathrm{E}[\mathrm{Var}(\boldsymbol{N}_T|\boldsymbol{\lambda})] = \phi \mathrm{E}[\boldsymbol{N}_T], \tag{11}$$

which is suggested to be based on the renewal theory, although the original paper does not specify the underlying assumptions about the spike generating process [5]. To see whether this equation might be consistent with the renewal theory, let us consider a renewal point process $\{g(\cdot), \lambda\}$ with a deterministic firing rate (constant in time and across trials), for which it holds [1, 2]

$$\lim_{T\to\infty} \frac{\mathrm{Var}(\boldsymbol{N}_T)}{\mathrm{E}[\boldsymbol{N}_T]} = \text{constant}.$$

However, this relation holds only in the limit of an infinite bin size $T \to \infty$ and for a constant firing rate. Before we can evaluate the relationship between $\mathrm{E}[\mathrm{Var}(\boldsymbol{N}_T|\boldsymbol{\lambda})]$ and $\mathrm{E}[\boldsymbol{N}_T]$ for a doubly stochastic renewal point process, we first need to define what this process is, that is, specify the spike generating process because in the absence of a mathematical model, the partitioning of variability is ambiguous [6]. Using our definition of a generative model for a doubly stochastic renewal point process, we derived the expression for $\mathrm{E}[\mathrm{Var}(\boldsymbol{N}_T|\boldsymbol{\lambda})]$ (Eq. 10 in Methods, Theorem 3), thus we can compute

$$\lim_{T\to\infty} \frac{\mathrm{E}[\mathrm{Var}(\boldsymbol{N}_T|\boldsymbol{\lambda})]}{\mathrm{E}[\boldsymbol{N}_T]} = \lim_{T\to\infty} \frac{\phi\mathrm{E}[\boldsymbol{N}_T] + \frac{1}{6} + \frac{1}{2}\phi^2 - \frac{1}{3}\psi(\phi) + \mathcal{O}(T^{-1})}{\mathrm{E}[\boldsymbol{N}_T]} = \phi.$$

Thus, the suggested relation Eq. 11 holds for our definition of a doubly stochastic point process, but also only for an infinite bin size in the limit $T \to \infty$. For a finite bin size, this ratio generally depends on the bin size because the expected spike count $\mathrm{E}[\boldsymbol{N}_T]$ increases with $T$. Therefore, the assumption Eq. 11 in the MR method is not usually true. It is valid for any bin size only in a special case when $\psi(\phi) = \frac{3}{2}\phi^2 + \frac{1}{2}$ which does not generally hold. For example, if the ISI probability density $g(\cdot)$ is the gamma distribution, then $\psi(\phi) = 2\phi^2$ (Methods), from which we see that this condition only holds when $\phi = 1$, that is, for the Poisson process.

Using the assumption Eq. 11, the MR method partitions the variability as

$$\mathrm{Var}(\mathrm{E}[\boldsymbol{N}_T|\boldsymbol{\lambda}]) = \mathrm{Var}(\boldsymbol{N}_T) - \phi\mathrm{E}[\boldsymbol{N}_T]. \tag{12}$$

Since variance must be non-negative ($\mathrm{Var}(\mathrm{E}[\boldsymbol{N}_T|\boldsymbol{\lambda}]) \geqslant 0$), the MR method estimates $\phi$ by calculating the minimum Fano factor estimated from data for all time bins:

$$\phi \approx \phi_{\mathrm{MR}} = \min_t \left\{ \frac{\mathrm{Var}(\boldsymbol{N}_T(t))}{\mathrm{E}[\boldsymbol{N}_T(t)]} \right\}. \tag{13}$$

This MR method for estimating $\phi$ contains multiple sources of bias. First, assuming the firing rate $\boldsymbol{\lambda}(t)$ changes smoothly on a timescale longer than the bin size $T$, we derived the correct partitioning equation for doubly stochastic renewal point processes (Eq. 10 in Methods, Theorem 3):

$$\mathrm{Var}(\mathrm{E}[\boldsymbol{N}_T|\boldsymbol{\lambda}]) = \mathrm{Var}(\boldsymbol{N}_T(t)) - \phi\mathrm{E}[\boldsymbol{N}_T(t)] - \frac{1}{6} - \frac{1}{2}\phi^2 + \frac{1}{3}\psi(\phi) + \mathcal{O}(T^{-1}).$$

Using the fact that $\mathrm{E}[\boldsymbol{N}_T(t)] = TE[\boldsymbol{\lambda}(t)]$ (Theorem 2) and $\mathrm{Var}(\boldsymbol{\lambda}(t)T) = T^2\mathrm{Var}(\boldsymbol{\lambda}(t))$, we can approximate the ratio of $\mathrm{Var}(\boldsymbol{N}_T(t))$ to $\mathrm{E}[\boldsymbol{N}_T(t)]$ for a finite bin size as:

$$\frac{\mathrm{Var}(\boldsymbol{N}_T(t))}{\mathrm{E}[\boldsymbol{N}_T(t)]} \approx \phi + T \cdot \frac{\mathrm{Var}(\boldsymbol{\lambda}(t))}{\mathrm{E}[\boldsymbol{\lambda}(t)]} + \frac{1}{T} \cdot \frac{\frac{1}{6} + \frac{1}{2}\phi^2 - \frac{1}{3}\psi(\phi)}{\mathrm{E}[\boldsymbol{\lambda}(t)]}.$$

From this equation and the definition of $\phi_{\mathrm{MR}}$ Eq. 13, we obtain the relation between the ground truth $\phi$ and $\phi_{\mathrm{MR}}$ estimated with the MR method:

$$\phi_{\mathrm{MR}} \approx \min_t \left\{ \phi + T \cdot \frac{\mathrm{Var}(\boldsymbol{\lambda}(t))}{\mathrm{E}[\boldsymbol{\lambda}(t)]} + \frac{1}{T} \cdot \frac{\frac{1}{6} + \frac{1}{2}\phi^2 - \frac{1}{3}\psi(\phi)}{\mathrm{E}[\boldsymbol{\lambda}(t)]} \right\}.$$

This equation shows that the MR method has several sources of bias. The first source is the term

$$T \cdot \frac{\mathrm{Var}(\boldsymbol{\lambda}(t))}{\mathrm{E}[\boldsymbol{\lambda}(t)]},$$

which is proportional to the trial-to-trial variance of the firing rate and linearly grows with the bin size $T$. This source of bias leads to an overestimation of $\phi$ and vanishes only when the variance of the firing rate is zero. The second source is the term

$$\frac{1}{T} \cdot \frac{\frac{1}{6} + \frac{1}{2}\phi^2 - \frac{1}{3}\psi(\phi)}{\mathrm{E}[\boldsymbol{\lambda}(t)]}.$$

We can predict the sign of this term in the case when $g(\cdot)$ is a gamma distribution. The sign is positive leading to an overestimation of $\phi$ if the ground-truth $\phi < 1$ (sub-Poisson spiking irregularity). The sign is negative leading to underestimation of $\phi$ if the ground-truth $\phi > 1$ (super-Poisson spiking irregularity). This source of bias decreases with increasing bin size and average firing rate. The third source of bias arises from the $\min_t\{\cdot\}$ operator. The term inside this operator is estimated for every bin from a finite number of trials in the data. Due to the estimation error, the estimated term in every bin comes from a sampling distribution with a finite width, which goes to zero as the number of trails goes to infinity. Assuming $\phi$ is constant across time bins [5], these estimates are samples drawn from the same distribution and the number of samples is equal to the number of time bins. Instead of reducing estimation noise by averaging samples, the operator $\min_t\{\cdot\}$ always chooses the minimum across samples, which always leads to underestimation of $\phi$, and this bias does not decrease with the number of samples.

Mixing these three sources of bias leads to unpredictable errors in $\phi_{\text{MR}}$ estimated with the MR method. $\phi_{\text{MR}}$ estimated by this method has been used to compute the variance of the firing rate $\text{Var}(\boldsymbol{\lambda}(t))$ using Eq. 13 [5], in which case the error in estimating $\phi$ propagates to the estimate of the firing-rate variance and how it depends on time. For example, if the mean firing rate increases with time, an underestimation of $\phi$ creates an increasing trend in $\text{Var}(\boldsymbol{\lambda}(t))$, even if the actual firing-rate variance is constant in time. Since $\text{Var}(\boldsymbol{\lambda}(t))$ is widely used as a metric to distinguish different classes of models for neural computation [5, 7, 8], unpredictable errors in the computation can lead to unreliable results and inaccurate interpretation of the data.

#### 1.4 Demšar comparison test for spiking irregularity $\phi$ across task conditions

| Condition | MR | MD | MAD | CI | Effect size | Magnitude |
|---|---|---|---|---|---|---|
| Cue-orth-2 | 2.316 | 1.153 | 0.398 | [1.062, 1.270] | 0.000 | negligible |
| Cue-orth-1 | 2.422 | 1.144 | 0.424 | [1.070, 1.274] | 0.024 | negligible |
| Cue-opp | 2.570 | 1.150 | 0.375 | [1.062, 1.275] | 0.010 | negligible |
| Cue-RF | 2.692 | 1.129 | 0.404 | [1.052, 1.245] | 0.062 | negligible |

**Table 1.** Demšar comparison test for $\phi$ between attention conditions in V4 data. MR - mean rank, MD - median, MAD - median absolute deviation, CI - confidence interval of MD, effect size - Cohen's effect size.

**Comparison of $\phi$ across attention conditions in V4.** We conducted the statistical analysis of the spiking irregularity $\phi$ for a population of 237 neurons in 4 attention conditions (c1: Cue-orth-2, c2: Cue-orth-1, c3: Cue-opp, c4: Cue-RF). The family-wise significance level of the tests is $\alpha = 0.05$. Based on the Shapiro-Wilk test, we rejected the null hypothesis that the population is normal for all conditions ($p < 10^{-4}$). Because we have more than two populations and all of them are not normal, we use the non-parametric Friedman test as an omnibus test to determine whether there are any significant differences between the median values of the populations. We use the post-hoc Nemenyi test to infer which differences are significant.

We report the median (MD), the median absolute deviation (MAD), and the mean rank (MR) among all populations over the samples (Table 1). Differences between populations are significant if the difference in the mean rank is greater than the critical distance CD=0.305 of the Nemenyi test. Based on the post-hoc Nemenyi test, we conclude that there are no significant differences within c1, c3, and c4 ($p = 0.071$). We also conclude that there are no significant differences within c1, c2, and c4 ($p = 0.106$). If we consider all four conditions together they are significantly different ($p = 0.009$) with the maximum effect size of 0.062 (Table 1). Since the effect size is smaller than the critical value of 0.2, it classifies as a negligible effect [9]. Therefore, we conclude that the differences in $\phi$ between the four attention conditions are negligible.

| Task | MR | MD | MAD | CI | Effect size | Magnitude |
|---|---|---|---|---|---|---|
| 4 choices | 1.483 | 0.546 | 0.291 | [0.448, 0.859] | 0.00 | negligible |
| 2 choices | 1.517 | 0.509 | 0.264 | [0.450, 0.788] | 0.14 | negligible |

**Table 2.** Demšar comparison test for $\phi$ during the two-choice and four-choice decision-making tasks in the LIP data. MR - mean rank, MD - median, MAD - median absolute deviation, CI - confidence of MD interval, effect size - Cohen's effect size.

**Comparison of $\phi$ between two-choice and four-choice decision-making tasks in LIP.** We conducted the statistical analysis of the spiking irregularity $\phi$ for a population of 60 neurons during four-choice (c1) and two-choice (c2) decision-making tasks. The family-wise significance level of the tests is $\alpha = 0.05$. Based on the Shapiro-Wilk test, we rejected the null hypothesis that the population is normal for both populations ($p < 10^{-4}$). No check for homogeneity was required because we only have two populations. Because we have only two populations and both of them are not normal, we use the Wilcoxon signed rank test to determine the differences in the central tendency and report the median (MD), the median absolute deviation (MAD), and the mean rank (MR) for each population (Table 2). We failed to reject the null hypothesis ($p = 0.109$) of the Wilcoxon signed rank test that population c2 (MD= $0.546 \pm 0.206$, MAD=0.291) is not greater than population c1 (MD= $0.509 \pm 0.169$, MAD=0.264). Therefore, we conclude that there is no statistically significant difference in $\phi$ between two-choice and four-choice decision-making tasks.

| Task epoch | MR | MD | MAD | CI | Effect size | Magnitude |
|---|---|---|---|---|---|---|
| Decision | 1.422 | 0.453 | 0.180 | [0.407, 0.735] | 0.00 | negligible |
| Fixation | 1.578 | 0.497 | 0.249 | [0.375, 0.753] | -0.204262 | small |

**Table 3.** Demšar comparison test for $\phi$ in the decision and fixation epochs of the decision-making task in LIP. MR - mean rank, MD - median, MAD - median absolute deviation, CI - confidence interval of MD, effect size - Cohen's effect size.

**Comparison of $\phi$ between decision and fixation epochs of the task in LIP.** We conducted the statistical analysis of the spiking irregularity $\phi$ for a population of 45 neurons during the decision (c1) and fixation (c2) epochs of the decision-making task. The family-wise significance level of the tests is $\alpha = 0.05$. Based on the Shapiro-Wilk test, we rejected the null hypothesis that the population is normal for the populations c1 ($p < 10^{-4}$) and c2 ($p = 0.005$). No check for homogeneity was required because we only have two populations. Because we have only two populations and both of them are not normal, we use the Wilcoxon signed rank test to determine the differences in the central tendency and report the median (MD), the median absolute deviation (MAD), and the mean rank (MR) for each population (Table 3). We failed to reject the null hypothesis ($p = 0.844$) of the Wilcoxon signed rank test that population c2 (MD= $0.453 \pm 0.164$, MAD=0.180) is not greater than population c1 (MD= $0.497 \pm 0.189$, MAD=0.249). Therefore, we conclude that there is no statistically significant difference in $\phi$ between decision and fixation epochs of the decision-making task in LIP.

**Comparison of $\phi$ across task conditions in PMd.** We conducted the statistical analysis of the spiking irregularity $\phi$ for a population of 272 neurons across 14 task conditions (c1 to c14,

| Task condition | MR | MD | MAD | CI | Effect size | Magnitude |
|---|---|---|---|---|---|---|
| c1 | 7.581 | 0.569 | 0.244 | [0.483, 0.661] | -0.05 | negligible |
| c2 | 7.618 | 0.579 | 0.252 | [0.488, 0.673] | -0.08 | negligible |
| c3 | 7.474 | 0.569 | 0.239 | [0.499, 0.664] | -0.05 | negligible |
| c4 | 7.346 | 0.576 | 0.278 | [0.498, 0.667] | -0.07 | negligible |
| c5 | 7.772 | 0.576 | 0.262 | [0.488, 0.650] | -0.07 | negligible |
| c6 | 7.305 | 0.558 | 0.238 | [0.503, 0.641] | 0.00 | negligible |
| c7 | 7.820 | 0.555 | 0.235 | [0.488, 0.652] | 0.01 | negligible |
| c8 | 8.033 | 0.550 | 0.258 | [0.472, 0.623] | 0.02 | negligible |
| c9 | 7.846 | 0.555 | 0.259 | [0.481, 0.641] | 0.01 | negligible |
| c10 | 7.522 | 0.570 | 0.260 | [0.490, 0.654] | -0.05 | negligible |
| c11 | 7.460 | 0.550 | 0.254 | [0.491, 0.656] | 0.03 | negligible |
| c12 | 6.967 | 0.557 | 0.276 | [0.492, 0.657] | 0.00 | negligible |
| c13 | 7.232 | 0.552 | 0.265 | [0.504, 0.662] | -0.02 | negligible |
| c14 | 7.026 | 0.574 | 0.244 | [0.487, 0.670] | -0.06 | negligible |

**Table 4.** Demšar comparison test of $\phi$ across task conditions in PMd. Task conditions (c1 to c14) correspond to seven stimulus coherence levels for each left and right response side. MR - mean rank, MD - median, MAD - median absolute deviation, CI - confidence interval of MD, and effect size - Cohen's effect size.

seven stimulus coherence levels for each left and right response side). The family-wise significance level of the tests is alpha=0.050. Based on the Shapiro-Wilk test, we rejected the null hypothesis that the population is normal for all 14 populations ($p < 10^{-4}$). Because we have more than two populations and all of them are not normal, we use the non-parametric Friedman test as an omnibus test to determine if there are any significant differences between the median values of the populations. We use the post-hoc Nemenyi test to infer which differences are significant. We report the median (MD), the median absolute deviation (MAD), and the mean rank (MR) among all populations over the samples (Table 4). Differences between populations are significant if the difference in the mean rank is greater than the critical distance CD=1.203 of the Nemenyi test. We failed to reject the null hypothesis ($p = 0.112$) of the Friedman test that there is no difference in the central tendency of the populations. Therefore, we conclude that there is no statistically significant difference between the median values of the populations.

| Task epoch | MR | MD | MAD | CI | Effect size | Magnitude |
|---|---|---|---|---|---|---|
| Decision | 1.710 | 0.461 | 0.183 | [0.403, 0.543] | 0.000 | negligible |
| Fixation | 1.290 | 0.530 | 0.159 | [0.468, 0.616] | -0.271 | small |

**Table 5.** Demšar comparison test for $\phi$ between the decision and fixation epochs of the decision-making task in PMd. MR - mean rank, MD - median, MAD: median absolute deviation, CI - confidence interval of MD, and effect size - Cohen's effect size.

**Comparison of $\phi$ between decision and fixation epochs of the trial in PMd.** We conducted the statistical analysis of the spiking irregularity $\phi$ for a population of 262 neurons during the decision (c1) and fixation (c2) epochs of the decision-making task. The family-wise significance level of the tests is alpha=0.050. Based on the Shapiro-Wilk test, we rejected the null hypothesis that the population is normal for the populations c1 ($p = 0.011$) and c2 ($p < 10^{-4}$). No check for homogeneity was required because we only have two populations. Because we have only two populations and both of them are not normal, we use the Wilcoxon's signed rank test to determine the differences in the central tendency and report the median (MD), the median absolute deviation (MAD), and the mean rank (MR) for each population (Table 5). We reject the null hypothesis ($p < 10^{-10}$) of the Wilcoxon's signed rank test that population c1 (MD= $0.461 \pm 0.070$, MAD=0.183) is not greater than population c2 (MD= $0.530 \pm 0.074$, MAD=0.159). Therefore, we conclude that the median of $\phi$ is significantly greater for c1 than c2 with an effect size of $-0.271$. This effect size is classified as a small effect size [9].
